# Elevated anti-SARS-CoV-2 antibodies and IL-6, IL-8, MIP-1β, early predictors of severe COVID-19

**DOI:** 10.1101/2021.04.13.439586

**Authors:** Helena Codina, Irene Vieitez, Alicia Gutierrez-Valencia, Vasso Skouridou, Cristina Martínez, Lucía Patiño, Mariluz Botero-Gallego, María Trujillo-Rodríguez, Ana Serna-Gallego, Esperanza Muñoz-Muela, María M. Bobillo, Alexandre Pérez, Jorge Julio Cabrera-Alvar, Manuel Crespo, Ciara K. O’Sullivan, Ezequiel Ruiz-Mateos, Eva Poveda, on behalf of the Cohort COVID-19 of the Galicia Sur Health Research Institute.

## Abstract

**Background:** Viral and host immune kinetics during acute COVID-19 and after remission of acute symptoms need better characterization.

**Methods:** SARS-CoV-2 RNA, anti-SARS-CoV-2 IgA, IgM, and IgG antibodies, and pro-inflammatory cytokines were measured in sequential samples among hospitalized COVID-19 patients during acute infection and 6 months following diagnosis.

**Results:** 24 laboratory-confirmed COVID-19 patients with mild/moderate and severe COVID-19 were included. Most were males 83%, median age of 61 years. 21% were admitted to the ICU and 8 of them (33.3%) met criteria for severe COVID-19 disease. A delay in SARS-CoV-2 levels decline during the first 6 days of follow-up and viral load persistence until month 3 were related with severe COVID-19, but not viral load levels at the diagnosis. Higher levels of anti-SARS-CoV-2 IgA, IgM, IgG and the cytokines IL-6, IL-8 and MIP-1β at the diagnosis time were related with severe COVID-19 outcome. Higher levels of MIP-1β, IL-1β, MIP-1*α* and IFN-*γ* were observed at month 1/3 during mild/moderate disease compared to severe COVID-19. IgG persisted at low levels after 6 months of diagnosis.

**Conclusions:** Higher concentrations of IgA, IgM, and IgG, and IL-6, IL-8 and MIP-1β are identified as early predictors of COVID-19 severity, but not SARS-CoV-2 RNA levels at diagnosis.

## INTRODUCTION

Coronavirus disease 2019 (COVID-19) caused by severe acute respiratory syndrome coronavirus 2 (SARS-CoV-2), rapidly spread worldwide becoming a global public health emergency. COVID-19 can be asymptomatic or mild in most cases, but it can rapidly progress to a severe lung inflammation leading to Acute Respiratory Distress Syndrome (ARDS), especially for older adults (> 80 years) and/or with comorbidities (i.e., serious heart conditions, chronic pulmonary disease, diabetes mellitus, or hypertension among others) [1–3]. The disease outcome mainly depends on the characteristics of the viral replication and host immune responses, which can end up resolving the infection efficiently or creating an exacerbated inflammation associated with severe lung damage pathology, organ failure and poor outcomes [4,5].

The comprehensive long-term kinetics of SARS-CoV-2 RNA levels, anti-SARS-CoV-2 antibodies (i.e., IgA, IgM, and IgG) and specific pro-inflammatory cytokines during and after COVID-19 is not fully characterized, and it is of great interest for the identification of early predictive biomarkers of severe disease and to understand the clinical outcomes of patients after the remission of acute symptoms.

This study comprises an evaluation of clinical, virological and immunological responses in a well-characterized cohort of hospitalized COVID-19 patients with mild/moderate and severe disease. We performed a close follow-up at different time points from the time of diagnosis and up to 6 months after the confirmation of the SARS-CoV-2 infection. Moreover, we were able to identify viral and host-immune biomarkers associated with COVID-19 severity. A delay in the SARS-CoV-2-RNA clearance in the upper respiratory tract during the first days of the disease, higher concentrations of anti-SARS-CoV-2 IgA, IgM, and IgG, and of specific cytokines (i.e., IL-6, IL-8 and MIP-1β) at baseline were associated with COVID-19 severity. Overall, after 6 months, anti-SARS-CoV-2 IgG persisted although at low levels.

## METHODS

### Patients, sample collection and clinical data

The study population was selected from the COVID-19 Cohort of the Galicia Sur Health Research Institute (COHVID-GS) (https://www.iisgaliciasur.es/apoyo-a-la-investigacion/cohorte-covid19/). This cohort includes laboratory-confirmed SARS-CoV-2 patients in clinical follow-up at the Vigo Healthcare Area with epidemiological/clinical data and with a repository of biological samples (i.e., nasopharyngeal swabs, serum, plasma and peripheral blood mononuclear cells-PBMCs) stored at the Galicia Sur Health Research Institute Biobank. The epidemiological/clinical information was collected in a Case Report Form (CRF) specifically predesigned for the COHVID-GS. The cohort also includes a control group of uninfected individuals (anti-IgA, IgM and IgG SARS-CoV-2 negative). Written informed consent was obtained from each individual enrolled in the COHVID-GS. All the techniques were performed in BLS-2 conditions, according to the biosafety guidelines for handling and processing specimens associated with COVID-19 (https://www.cdc.gov/coronavirus/2019-ncov/lab/lab-biosafety-guidelines.html).

The selection criteria for this study were SARS-CoV-2 adult hospitalized patients with a very close clinical follow-up with nasopharyngeal swabs, serum and plasma samples, with consecutive samples available at least at baseline; and day 3 or 6; and month 1 or 3 from their inclusion in the COHVID-GS. Moreover, only those patients included in the COHVID-GS between day 1 and 5 following the confirmation of the SARS-CoV-2 infection by RT-qPCR were considered. Serial paired nasopharyngeal swabs, serum and plasma samples were collected from each patient at baseline, day 3 or 6, month 1 or 3 and at month 6. We also included 30 serum samples from healthy and non-infected individuals as controls for the immunoassay experiments. Epidemiological and clinical data were recorded for the study population.

Following WHO guidance, severe COVID-19 was defined as the need for invasive mechanical ventilation, the development of Acute Respiratory Distress Syndrome (PO2/FiO2 < 300 mmHg or Saturation O2/FiO2 < 330) or the admission to an Intensive Care Unit (ICU) [6]. Patients who did not meet severe criteria were considered as mild/moderate COVID-19.

### SARS-CoV-2 RNA extraction

Viral RNA was extracted from 140 µL of nasopharyngeal swab samples using the QIAmp viral RNA mini kit (QIAGEN, Hilden, Germany) and the automatized QIAcube system (QIAGEN, Hilden, Germany), and was eluted in 50 µL of buffer following the manufacturer’s instructions. The positive (SARS-CoV-2 Standard, EDx Exact Diagnostics) and negative (SARS-CoV-2 Negative, EDx Exact Diagnostics) controls were also extracted using the same procedure.

### Droplet digital PCR analysis

SARS-CoV-2 viral load was quantified by reverse transcriptase droplet digital PCR (RT-ddPCR), including the one-step reverse transcription (One-Step RT-ddPCR Advanced Kit for Probes, Bio-Rad Laboratories) and the triplex probe assay for PCR amplification (2019-nCoV CDC ddPCR Triplex Probe Assay, Bio-Rad Laboratories). The assay contains primers and probes targeting two regions of the SARS-CoV-2 nucleocapsid gene (N1 and N2) and the human RNase P gene (*RPP30*). The reaction mixture was performed with 5.5 µL of SARS-CoV-2 RNA sample and following the manufacturer’s instructions. All the samples were tested in duplicate. Data analysis was performed using the QuantaSoft Analysis Pro Software (Bio-Rad Laboratories) which showed the results as copies per microliter of 1x ddPCR reaction. All viral load values were recalculated to copies per millilitre of swab, and the sensitivity threshold was 100 copies/mL. To assess the accuracy of the absolute viral RNA quantification, two-fold serial dilutions of the positive control were analysed for lineal regression analysis. Finally, to establish a reliable comparison between the viral load of different samples, the absolute quantification of SARS-CoV-2 RNA was normalized with *RPP30* and expressed as copies/10^4^ cells (supplementary material section).

### Anti-SARS-CoV-2 IgA, IgG, and IgM quantification

The wells of 96-well immunoassay plates were coated overnight at 4°C with 50 μL of 5 μg/mL of SARS-CoV-2 nucleoprotein (NP) in 50 mM carbonate buffer pH 9.4. The wells were washed three times with 200 μL of PBS containing 0.05 % (v/v) Tween-20 (PBST) and then blocked with 200 μL of 5 % (w/v) skim milk in PBST for 30 min. After another washing step, 50 μL of serum samples (diluted 1/100 with PBS after heating for 30 min at 56°C to inactivate residual virus) were added to the wells and incubated for 1 h. The wells were washed again three times with PBST and 50 μL of anti-human IgA-HRP, anti-human IgM-HRP or anti-human IgG-HRP enzyme conjugates diluted 1/20000 with PBST were added. After a final incubation for 30 min, the wells were washed five times with PBST and 50 μL of TMB Super Sensitive ELISA substrate were added in each well. The reaction was stopped by the addition of 50 μL of 1 M H2SO4 after 5 min for IgM and IgA detection or 7 min for IgG detection. The absorbance at 450 nm was finally read on a SPECTRAmax 340PC-384 microplate reader. The levels of IgA, IgM and IgG antibodies in each sample were estimated using standard antibody calibration curves performed in parallel in each plate as detailed in the supplementary information. Pre-pandemic serum samples were used as controls to set the background levels of the in-house developed ELISA. All incubation steps were performed at room temperature (22-25°C) unless stated otherwise.

### Plasma cytokines quantification

Plasma cytokines concentrations were determined using enzyme-linked immunosorbent assays. Interleukin-6 (IL-6), interleukin-8 (IL-8), interleukin 1 beta (IL-1β), Tumor Necrosis Factor-alpha (TNF-α), interferon-gamma (IFN-γ), Macrophage Inflammatory Proteins 1 alpha (MIP-1α) and 1 beta (MIP-1β) were analysed using a multiplex bead-based immunoassay (MILLIPLEX® MAP Human High Sensitivity T Cell Magnetic Bead Panel). Interferon gamma-induced protein 10 (IP-10), (Human CXCL10 ELISA kit, Abcam, Cambridge, UK) and soluble receptor interleukin 2 (sCD25) were analysed using the Human Quantikine Immunoassay (R&D Systems). All following the manufacturer’s instructions.

### Statistics

The descriptive analyses were reported as frequencies and percentages for categorical variables and medians and interquartile ranges (IQRs) for continuous variables. To compare antibody levels between COVID-19 patients and a peer control group, and to compare SARS-CoV-2 viral load and cytokines between severe and mild/moderate patients, Mann-Whitney U test was performed. To assess the differences in the viral load overtime and the concentration of the cytokines in several time points during the first 6 months after SARS-COV-2 infection, a Wilcoxon Signed Rank test was performed. Finally, Mann-Whitney U test was also used to compare the baseline antibodies levels and baseline cytokines concentration with the severe clinical outcomes (ICU admission, requirement of invasive mechanical ventilation and development of Acute Respiratory Distress Syndrome). Statistical analyses were done with SPSS version 19 and GraphPad Prism 8.2.1 and a p value less than 0.05 was considered statistical significance.

## RESULTS

### Demographic and clinical characteristics of the studied patients

A total of 24 hospitalized laboratory-confirmed COVID-19 patients were included in the study. The demographic and most relevant clinical characteristics related with COVID-19 are shown in **Table 1**. Most 83% (20/24) were males with a median age of 61 years old (IQR: 48-75.3). Nearly 50% (12/24) of patients met obese criteria (BMI ≥ 30) and 42% (10/24) had hypertension. The median hospitalization time was 8 days, and 21% (5/24) were admitted to the ICU. Eight of them (33.3%) met criteria for severe COVID-19 disease.

**Table 1.** Demographic and clinical characteristics of the study population.

| Characteristics | n=24 |
| --- | --- |
| Age (years) | 61.0 (48.0 – 75.3) |
| Sex (male) | 83% (20) |
| Comorbidities |  |
| Obesity BMI>30 | 50% (12) |
| Hypertension | 42% (10) |
| Chronic lung disease | 21% (5) |
| Dyslipidemia | 21% (5) |
| Cardiovascular disease | 13% (3) |
| Diabetes | 13% (3) |
| Hospitalization time (days) | 8.0 (7.0 – 16.3) |
| ICU admission | 21% (5) |
| ICU length of stay (days) | 12.0 (9.0 – 18.5) |
| Exitus | 4% (1) |
| Time from symptom onset to hospital admission (days) | 9.0 (5.5 – 13.0) |
| Time from hospital admission to cohort admission (days) | 2.0 (2.0 – 3.0) |
| Time from PCR+ to cohort admission (days) | 3.0 (2.0 – 5.0) |
| Symptoms on admission |  |
| Fever | 75% (18) |
| Dyspnea | 71% (17) |
| Cough | 67% (16) |
| Malaise | 67% (16) |
| Diarrhea | 42% (10) |
| Sputum | 38% (9) |
| ARDS* | 30% (7) |
| Myalgia/Arthralgia | 30% (7) |
| Anosmia | 25% (6) |
| Chest pain | 25% (6) |
| Median laboratory values on admission |  |
| Leukocytes (cells/mL) | 4940.0 (4545.0 – 6815.0) |
| Lymphocytes (cells/mL) | 955.0 (755.0 – 1215.0) |
| Neutrophils (cells/mL) | 3725.0 (2860.0 – 5437.5) |
| Platelets (x10<sup>9</sup>/L) | 166.5 (132.8 – 248.0) |
| Creatinine (mg/dL) | 0.885 (0.778 – 1.023) |
| LDH (UI/L) ** | 274.0 (208.0 – 350.0) |
| Bilirubin (mg/dL)** | 0.660 (0.430 – 0.900) |
| Oxygen therapy | 58% (14) |
| Invasive mechanical ventilation | 21% (5) |
| Pulmonary infiltrates | 92% (22) |
| Treatments during hospitalization |  |
| Hydroxychloroquine | 71% (17) |
| Antibiotics | 46% (11) |
| Immunosuppressive therapy | 46% (11) |
| Corticosteroids | 42% (10) |
| Lopinavir/Ritonavir | 42% (10) |
| Azithromycin | 30% (7) |

Data are median (IQR), % (n).
\*Acute Respiratory Distress Syndrome.
\*\* n=23

### SARS-CoV-2 viral load kinetics

The SARS-CoV-2 viral load quantification was performed from 92 RNA samples extracted from nasopharyngeal swabs in the 24 patients at different time-points (24 at baseline, 18 at day 3, 13 at day 6, 5 at day 15, 16 at month 1, and 16 at month 3) (**Table 2)**. At baseline, 23 of 24 patients (96%) were positive for SARS-CoV-2 with a median viral load of 2227.17 copies/10^4^ cells showing broad variability between patients (IQR: 69.78 – 7.42×10^4^). Only one patient was negative for SARS-CoV-2 RNA quantification but with low amplification of *RPP30* indicating a deficient sampling. Overall, SARS-CoV-2 viral load decrease overtime, and most of the patients (80%) were negative one month after the diagnosis **(Figure 1A)**. However, 5 patients showed persistent viral load at month 1 and 3. All these patients had severe disease (COV 006, COV 009 and COV 013) and/or comorbidities such as diabetes (COV 016 and COV 013); hypertension or cardiovascular disorders (COV 016, COV 019 and COV 013); chronic lung disease and HIV infection (COV 009) and dyslipidemia (COV 006). Two different profiles in SARS-CoV-2 kinetics were observed during acute infection based on the COVID-19 clinical outcomes. A delay for the significant decline of SARS-CoV-2 levels during the first 6 days of follow-up was recognized for those patients with severe disease. Thus, while for patients with mild/moderate disease a significant decline in viral load is observed since the diagnosis time, for patients with severe disease is only observed after day 6 (**Figure 1B)**.

**Table 2.**
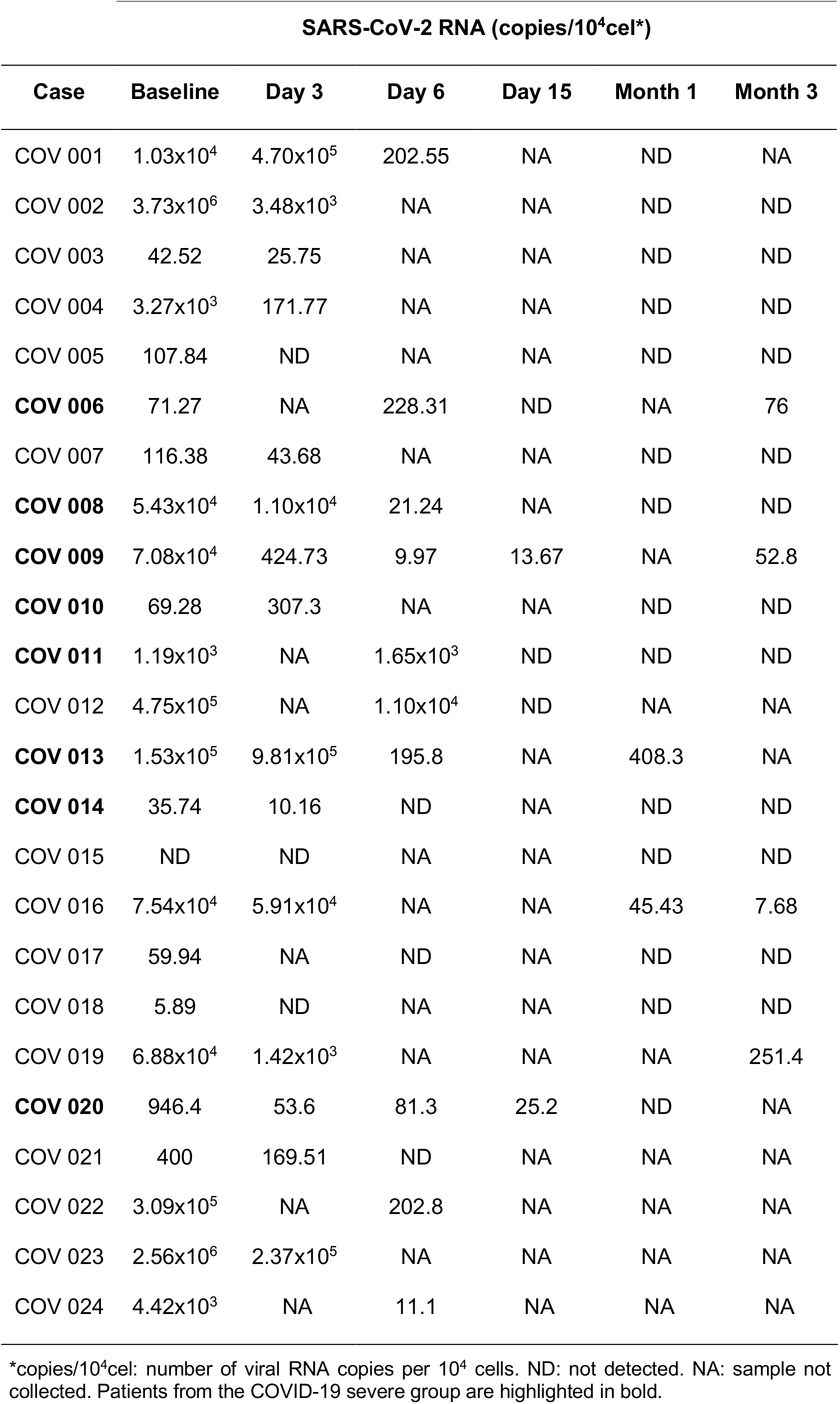
SARS-CoV-2 RNA quantification from nasopharyngeal swab samples.

| Case | SARS-CoV-2 RNA (copies/10<sup>4</sup>cel*) |  |  |  |  |  |
| --- | --- | --- | --- | --- | --- | --- |
|  | Baseline | Day 3 | Day 6 | Day 15 | Month 1 | Month 3 |
| COV 001 | 1.03x10<sup>4</sup> | 4.70x10<sup>5</sup> | 202.55 | NA | ND | NA |
| COV 002 | 3.73x10<sup>6</sup> | 3.48x10<sup>3</sup> | NA | NA | ND | ND |
| COV 003 | 42.52 | 25.75 | NA | NA | ND | ND |
| COV 004 | 3.27x10<sup>3</sup> | 171.77 | NA | NA | ND | ND |
| COV 005 | 107.84 | ND | NA | NA | ND | ND |
| **COV 006** | 71.27 | NA | 228.31 | ND | NA | 76 |
| COV 007 | 116.38 | 43.68 | NA | NA | ND | ND |
| **COV 008** | 5.43x10<sup>4</sup> | 1.10x10<sup>4</sup> | 21.24 | NA | ND | ND |
| **COV 009** | 7.08x10<sup>4</sup> | 424.73 | 9.97 | 13.67 | NA | 52.8 |
| **COV 010** | 69.28 | 307.3 | NA | NA | ND | ND |
| **COV 011** | 1.19x10<sup>3</sup> | NA | 1.65x10<sup>3</sup> | ND | ND | ND |
| COV 012 | 4.75x10<sup>5</sup> | NA | 1.10x10<sup>4</sup> | ND | NA | NA |
| **COV 013** | 1.53x10<sup>5</sup> | 9.81x10<sup>5</sup> | 195.8 | NA | 408.3 | NA |
| **COV 014** | 35.74 | 10.16 | ND | NA | ND | ND |
| COV 015 | ND | ND | NA | NA | ND | ND |
| COV 016 | 7.54x10<sup>4</sup> | 5.91x10<sup>4</sup> | NA | NA | 45.43 | 7.68 |
| COV 017 | 59.94 | NA | ND | NA | ND | ND |
| COV 018 | 5.89 | ND | NA | NA | ND | ND |
| COV 019 | 6.88x10<sup>4</sup> | 1.42x10<sup>3</sup> | NA | NA | NA | 251.4 |
| **COV 020** | 946.4 | 53.6 | 81.3 | 25.2 | ND | NA |
| COV 021 | 400 | 169.51 | ND | NA | NA | NA |
| COV 022 | 3.09x10<sup>5</sup> | NA | 202.8 | NA | NA | NA |
| COV 023 | 2.56x10<sup>6</sup> | 2.37x10<sup>5</sup> | NA | NA | NA | NA |
| COV 024 | 4.42x10<sup>3</sup> | NA | 11.1 | NA | NA | NA |

\*copies/10<sup>4</sup>cel: number of viral RNA copies per 10<sup>4</sup> cells. ND: not detected. NA: sample not collected. Patients from the COVID-19 severe group are highlighted in bold.

**Figure 1.**
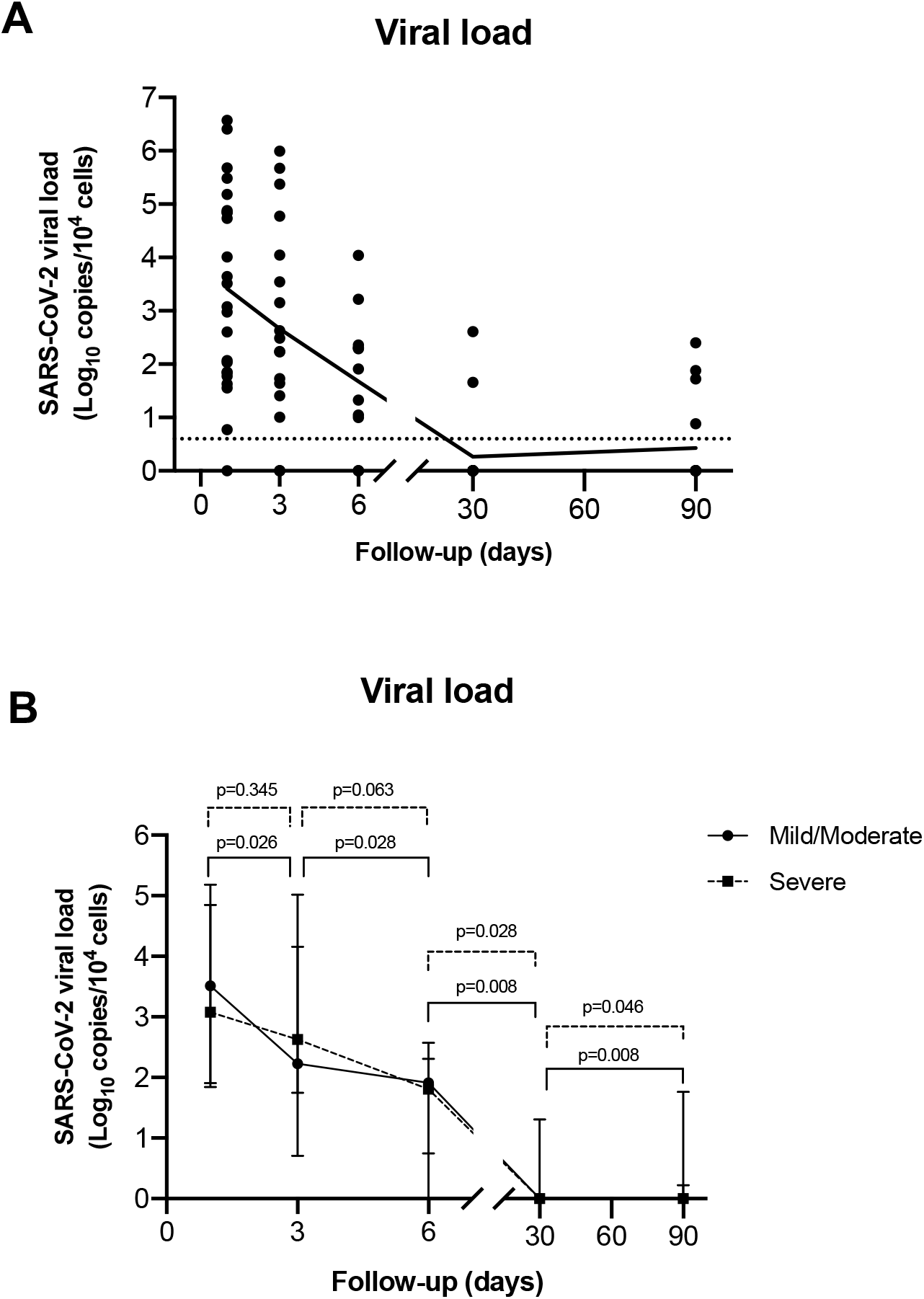
SARS-CoV-2 viral load kinetics. A) SARS-CoV-2 RNA levels among study period. B) Median SARS-CoV-2 RNA levels in patients with mild/moderate versus severe COVID-19. A) SARS-CoV-2 RNA levels among the overall study population during the study period. The black dots represent the aligned individual values obtained from each time-point. The solid line represents the connected mean values. The dotted line indicates the established experimental threshold (below the lowest positive sample obtained in our study). B) Median ± IQR SARS-CoV-2 RNA levels in patients with mild/moderate versus severe COVID-19. The solid line with black circles represents the mild/moderate group and dotted line with black squares represents the severe group. Wilcoxon Signed Rank Test was performed for the decline of viral load between time-points; p: p-value.

### Anti-SARS-CoV-2 antibodies kinetics

The anti-SARS-CoV-2 antibodies levels were quantified at different time-points from a total of 109 serum samples (24 at baseline, 18 at day 3, 13 at day 6, 5 at day 15, 16 at month 1, 18 at month 3 and 15 at month 6) (**Figure 2**). Overall, a high variability in the IgA baseline levels was observed among patients that progressively increase reaching the highest median levels at day 6 of follow-up, and then, significantly decrease until month 6 (p= 0.028). The highest levels of IgA at baseline and during the follow-up were observed for patients with severe COVID-19 (COV 006, COV 008, COV 010, COV 014, COV 013) (**Figure 2A**). A similar pattern was observed for the kinetics of IgM with an on-going increase reaching the maximum levels at day 6 with a significant decline after that (p=0.028). Likewise to IgA, the highest levels of IgM at baseline and during the follow-up were observed for patients with severe COVID-19 (COV 008, COV 011, COV 010, COV 020) (**Figure 2B**). For IgG there was also a gradual increase until day 6 with a progressive but slowly decrease until month 6 reaching very low concentrations (**Figure 2C**). Of note, IgG levels at month 6 were significantly higher than for a group of 30 uninfected controls (15.93 vs. 4.50 μg/mL, respectively p-value < 0.001) (**Figure 2D**).

**Figure 2.**
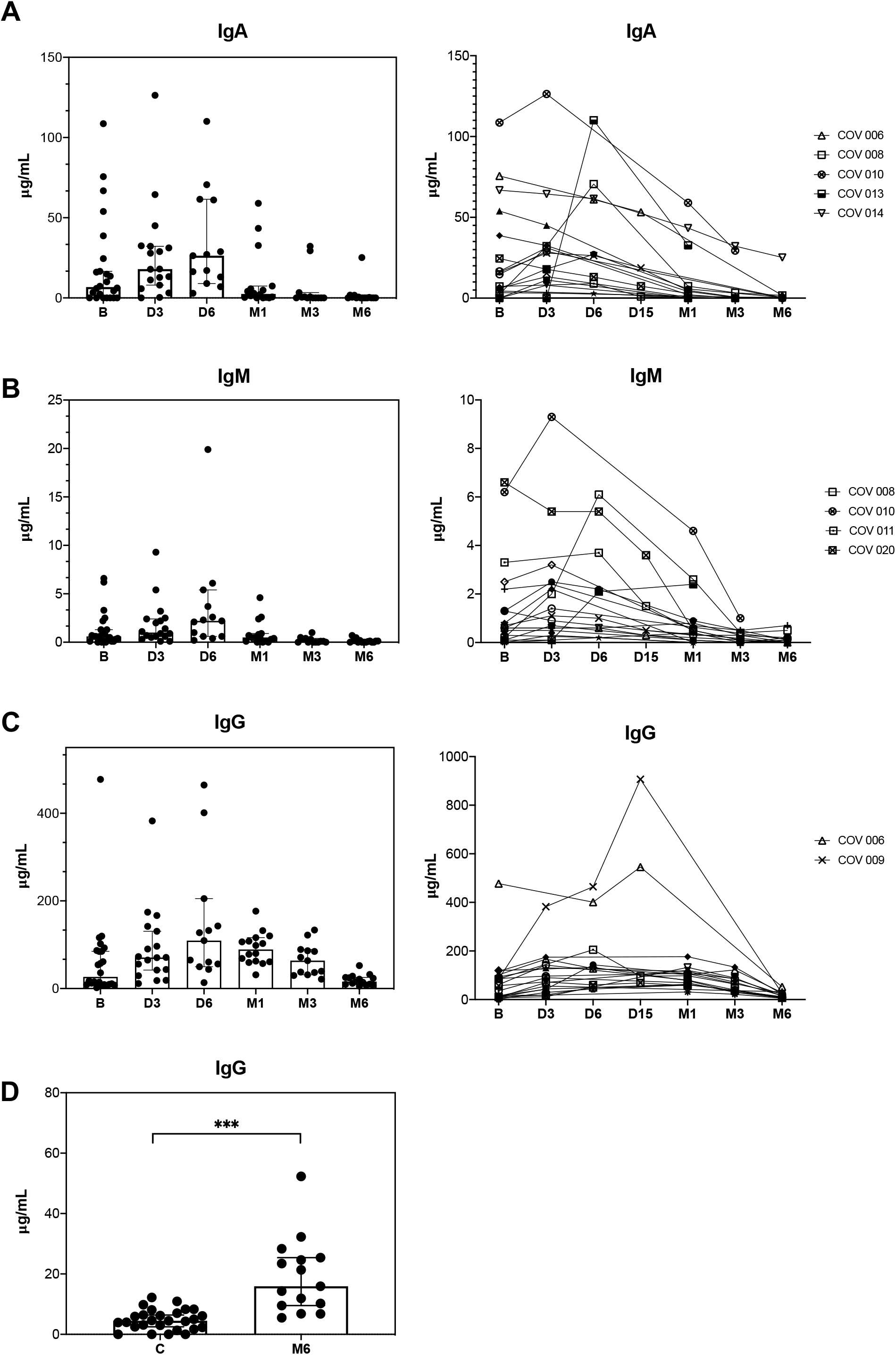
Absolute levels at different time points during the study period and antibodies IgA (A), IgM (B) and IgG (C) kinetics in specific patients. D) Comparison of IgG levels between uninfected controls and patients after 6 months of SARS-CoV-2 infection. Absolute levels of IgA (A), IgM (B) and IgG (C) among the overall study population (n=24) at baseline (B), day 3 (D3), day 6 (D6), month 1 (M1), month 3 (M3) and month (M6) after SARS-CoV-2 infection and changes of serum IgA (A), IgM (B) and IgG (C) antibodies in specific patients (n=19) at baseline (B), day 3 (D3), day 6 (D6), month 1 (M1), month 3 (M3) and month (M6) after SARS-CoV-2 infection. Patients COV 006, COV 008, COV 009, COV 010, COV 011, COV 013, COV 014 and COV 020 belong to the severe group. D) Difference in IgG levels between uninfected controls (n=30) and patients after 6 months of SARS-CoV-2 infection (n=15) was determined by Mann-Whitney U test. ***, p-value < 0.0001.

The numerical data for IgA, IgM and IgG in serum during the study period are shown as supplementary data (**Supplementary Table 1)**.

### Plasma cytokines kinetics

Plasma cytokines levels were performed in a total of 18 patients, 12 with mild/moderate and 6 with severe COVID-19, at different time-points (18 at baseline, 15 at month 1, 14 at month 3, and 17 at month 6). **Figure 3** shows the dynamic of the median concentrations of the 9 cytokines assessed (TNF-*α*, IL-6, IL-8, IL-1β, MIP-1*α*, MIP-1β, IFN-*γ*, sCD25 and IP-10) for each group based on COVID-19 severity during the study period.

**Figure 3.**
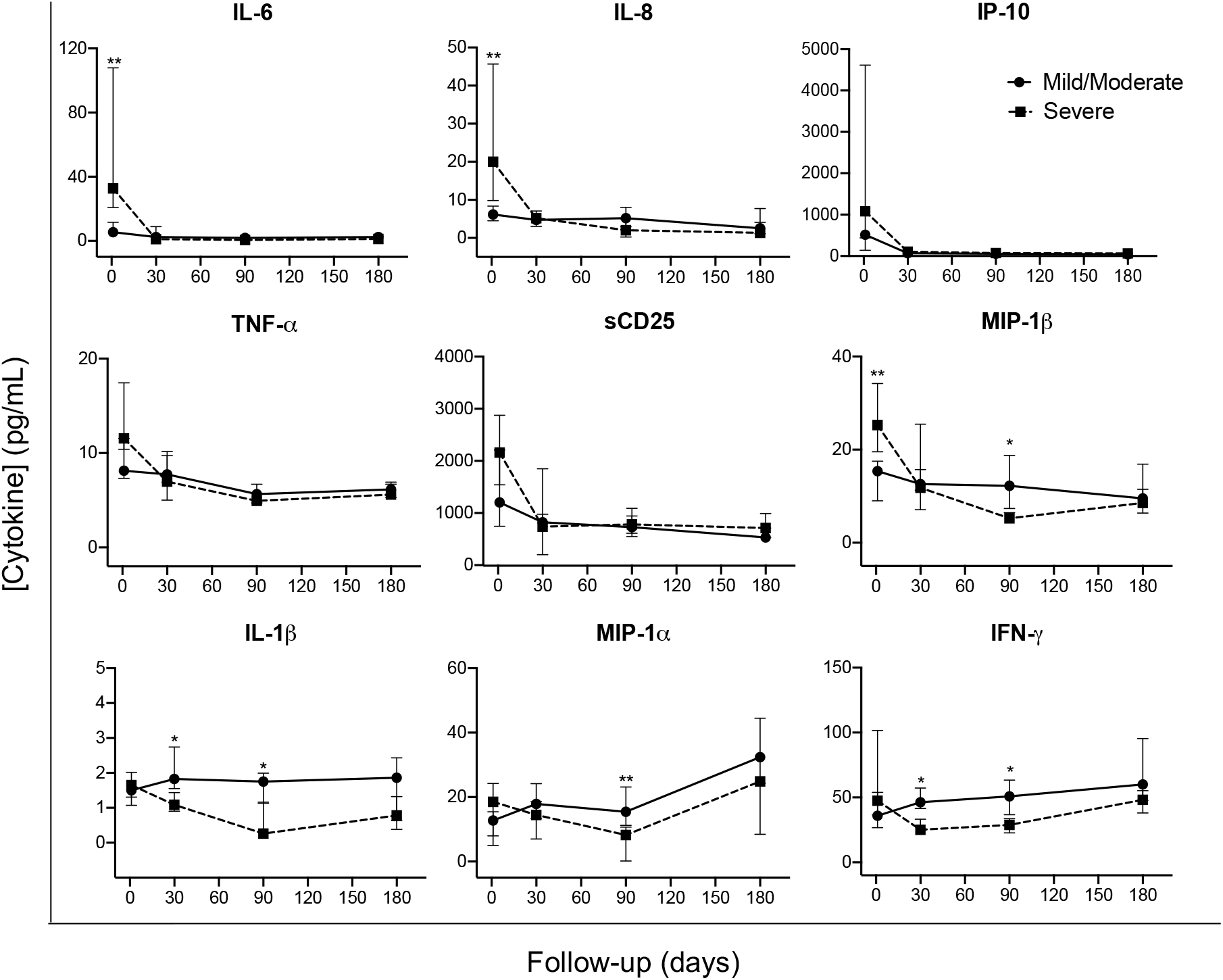
Cytokine kinetics in patients with mild/moderate versus severe COVID-19 during the study period. Cytokine levels are expressed as median ± IQR. The solid line with black circles represents the mild/moderate group (n=12) and dotted line with black squares represents severe group (n=6). Differences between both groups at each time-point was assessed using Mann-Whitney U test. *, p-value < 0,05; **, p-value < 0.001.

At baseline, significant higher levels of IL-6, IL-8, and MIP-1β were observed for patients with severe disease compared with those with mild/moderate COVID-19. Overall, a similar dynamic was observed for some of the pro-inflammatory markers, IL-6, IL-8, IP-10, TNF-*α* and sCD25 during the study period. By contrast, the dynamic for MIP-1β, IL-1β, MIP-1*α* and IFN-*γ* showed a different profile during the follow-up with significant higher levels at month 1 (IL-1β and IFN-*γ*) and month 3 (MIP-1β, IL-1β, MIP-1*α* and IFN-*γ*) among patients with mild/moderate diseases compared to those with severe COVID-19. After 6 months of the acute SARS-COV-2 infection no differences were found between patients with mild/moderate vs. severe disease.

The numerical data for plasma cytokines comparison between mild/moderate vs. severe patients are shown as supplementary data (**Supplementary Table 2)**.

### Clinical, virological and host immune biomarkers for severe COVID-19 outcomes

A univariate analysis was performed to identify potential biomarkers associated with severe clinical outcomes, defined as the requirement of invasive mechanical ventilation, the admission to an Intensive Care Unit (ICU) and the development of Acute Respiratory Distress Syndrome (ARDS). Higher levels of IgA, IgM, IgG against SARS-CoV-2 and also the cytokines IL-6, IL-8 and MIP-1β at baseline was observed among patients who developed ARDS compared to those without ARDS. Moreover, higher levels of IL-6 and MIP-1β were related with either the need of invasive mechanical ventilation or ICU admission. Overall, SARS-CoV-2 viral load at baseline was not associated with any of the severe clinical outcome definitions (**Table 3)**.

**Table 3.** Biomarkers of severe COVID-19.

|  | ARDS | No ARDS | p-value | Invasive mechanical ventilation/<br>ICU admission | No invasive mechanical ventilation/<br>No ICU admission | p-value |
| --- | --- | --- | --- | --- | --- | --- |
| SARS-CoV-2 viral load<br>(Log<sub>10</sub> copies/10<sup>4</sup> cells) | 2.97 (1.78 - 4.73) | 3.64 (1.92 - 5.36) | 0.357 | 3.07 (2.70- 5.01) | 3.51 (1.71- 4.88) | 0.804 |
| Serum antibodies<sup>a</sup> (µg/mL) |  |  |  |  |  |  |
| IgA | 24.51 (7.25 - 75.62) | 4.42 (0.0 - 14.32) | **0.013** | 6.18 (0 - 16.63) | 7.25 (1.22 - 50.07) | 0.783 |
| IgG | 84.94 (57.32 - 102.03) | 14.47 (8.38 - 57.71) | **0.028** | 0.56 (0.10 - 1.30) | 0.59 (0.17 - 4.96) | 0.446 |
| IgM | 0.66 (0.59 - 6.25) | 0.43 (0.09 - 1.06) | **0.047** | 17.42 (8.96 - 86.86) | 57.32 (6.47 - 281.21) | 0.836 |
| Plasma cytokines<sup>b</sup> (pg/mL) |  |  |  |  |  |  |
| IL-6 | 32.72 (20.78 - 107.92) | 5.44 (2.93 - 11.63) | **0.002** | 38.51 (32.72 - 172.42) | 5.80 (3.72 - 16.57) | **0.017** |
| IL-8 | 19.99 (9.83 - 45.67) | 6.43 (4.64 - 9.51) | **0.007** | 21.56 (15.78 - 51.42) | 7.69 (5.02 - 10.88) | 0.056 |
| IP-10 | 1083.80 (439.15 - 4613.25) | 515.40 (141.3 - 633.13) | 0.102 | 1606.00 (1076.40 - 3347.00) | 534.70 (177.65 - 614.15) | 0.100 |
| TNF-α | 11.55 (10.40 - 17.45) | 8.12 (7.32 - 11.56) | 0.083 | 10.96 (10.60 - 11.92) | 10.81 (7.55 - 11.73) | 0.574 |
| sCD25 | 2160.00 (1541.25 - 2877.0) | 1221.50 (775.33 - 2806.00) | 0.250 | 1567.00 (1515.00 - 2247.50) | 1465.00 (960.95-2534.50) | 0.645 |
| MIP-1β | 25.29 (19.52 - 34.19) | 15.37 (8.99 - 17.51) | **0.001** | 30.82 (24.47 - 37.55) | 16.60 (9.98 - 19.99) | **0.039** |
| IL-1β | 1.65 (1.07 - 1.77) | 1.50 (1.31 - 2.02) | 0.750 | 1.62 (1.42 - 1.73) | 1.60 (1.30 - 1.80) | 1.000 |
| MIP-1α | 18.52 (5.02 - 24.21) | 12.74 (7.95 - 15.47) | 0.682 | 15.11 (10.87 - 18.52) | 13.36 (8.34 - 19.62) | 1.000 |
| IFN-γ | 47.59 (36.60 - 101.63) | 35.91 (26.79 - 53.89) | 0.213 | 42.13 (40.57 - 77.18) | 37.01 (30.6 - 53.66) | 0.301 |

Data are median (IQR) Mann-Whitney U test was performed. P-values < 0.05 were considered significant and are highlighted in bold. a. ARDS n=7; No ARDS n=17; ICU admission n=5; No ICU admission n=19 b. ARDS n=6; No ARDS n=12; ICU admission n=3; No ICU admission n=15

## DISCUSSION

In this study we report that elevated levels of anti-SARS-CoV-2 antibodies, IgA, IgM, and IgG as well as specific cytokines such as IL-6, IL-8, MIP-1β during SARS-CoV-2 acute infection are associated with severe COVID-19, defined by the development of ARDS or the need of invasive mechanical ventilation or UCI admission. IgG persists after 6 months of the diagnosis but at lower concentrations. Whilst the levels of some cytokines (i.e., IL-6, IL-8, IP-10, TNF-*α*, sCD25) decline one month after of SARS-CoV-2 diagnosis, others (MIP-1β, IL-1β, MIP-1*α* and IFN-*γ*) showed higher levels at month 1 and/or 3 for patients with mild/moderate disease compared to those with severe COVID-19. However, after 6 months after diagnosis, no differences in cytokines levels between patients with mild/moderate vs. severe disease were observed. Interestingly, although SARS-CoV-2 RNA levels were not associated with the clinical outcome, during the acute infection, a significant delay in SARS-CoV-2 viral load decline was recognized for patients with severe COVID-19 as compared to those with mild/moderate COVID-19 during the first 6 days of follow-up.

Overall, SARS-CoV-2 viral load kinetics follows the pattern described previously, reaching maximum levels during acute infection and decreasing progressively until becoming undetectable in most patients in less than one month following diagnosis [7– 10]. However, because of the close follow-up of these patients we were able to identify a significant delay during the first days of viral load decay for patients who met the criteria for severe COVID-19. Nonetheless, in our cohort, SARS-CoV-2 RNA levels were not associated with the clinical outcome at any time point. There are some controversial data regarding this issue, while some studies have not reported any association between SARS-CoV-2 viral load during acute infection and COVID-19 severity [11,12] others point out that elevated viral load could be used to identify patients at higher risk for morbidity or severe COVID-19 outcome [13,14]. The controversial results might be explained for the heterogeneity of the studies related to the different characteristics of the study population and/or the methodology used for SARS-CoV-2 quantification and sampling quality (i.e., normalization using copies/cells).

Although, 80% of patients showed undetectable viremia one month after the diagnosis, in 5 patients we observed SARS-CoV-2 RNA persistence between month 1 and 3. Interestingly, all of these had severe disease and/or comorbidities (i.e., diabetes, hypertension or cardiovascular disorders, chronic lung disease and HIV infection, dyslipidemia). These observations are in agreement with previous studies that reported an association between SARS-CoV-2 RNA persistence with severe disease [9,15] and comorbidity [13,16–18] following remission of the acute symptoms. More recently, Jacobs et al., highlighted the association between SARS-CoV-2 plasma viremia with ICU admission [19] and Tokuyama et al., described SARS-CoV-2 persistence in intestinal enterocytes up to 7 months after symptoms resolutions [20]. Additional studies are required to better understand the clinical significance of SARS-CoV-2 persistence in different compartments (i.e., plasma, gut, upper respiratory tract) and its potential use to monitor COVID-19 patients during the early acute infection and/or following the remission of the acute symptoms.

We found that elevated levels of anti-SARS-CoV-2 antibodies during acute infection are related with severe COVID-19. Those patients who developed ARDS showed significantly higher levels of IgA, IgM, and IgG, compared to patients without ARDS. These findings are in agreement with previous studies that found an association between anti-SARS-CoV-2 antibodies levels and the need for intubation or death [21,22]. Overall, we observed that anti-SARS-CoV-2 IgA and IgM antibodies decay rapidly after the acute infection phase (i.e., at month 3, 54% for IgA and 77% for IgM showed levels below the limit of detection) but the decline for IgG was less prominent and persisted until month 6. These levels were still significantly higher compared to the levels of uninfected controls (15.93 vs. 4.50 μg/mL, respectively p-value < 0.001). The durability of the immune responses against SARS-CoV-2 infection is a matter of intense interest and still remains unclear. Regarding the humoral responses, recent studies have also reported that IgG levels persist between 6 and 8 months after onset of symptoms [23,24]. However, plasma neutralization activity seems to decrease a few weeks after the onset of the symptoms [25]. Understanding the dynamics of antibodies against SARS-CoV-2 and the persistence of the neutralizing activity is critical to establish correct prevention and vaccination strategies.

COVID-19 severity has been associated with an exacerbated inflammation due to a massive release of pro-inflammatory components [4,5]. We have identified that elevated levels of IL-6, IL-8, MIP-1β during SARS-CoV-2 acute infection are associated with severe COVID-19 defined by the development of ARDS or the need of invasive mechanical ventilation or UCI admission. Higher levels of IL-6 have been consistently related with severe COVID-19 disease [26,27] and play a pivotal role in the cytokine storm in response to SARS-CoV-2 infection promoting organ failure and severe lung pathology [28–30]. IL-8 leads the activation and the recruitment of neutrophils to the inflammation sites and has been implicated in inflammatory pulmonary diseases such as ARDS, chronic obstructive pulmonary disease, and asthma [31,32]. MIP-1β drives the recruitment of a variety of innate and adaptive immune cells and high concentrations have been reported in the serum and lungs of patients with certain acute respiratory viral infections [33,34]. However, their role during SARS-CoV-2 infection remains unclear. Some studies have reported a higher production of MIP-1β at the transcriptional level in bronchoalveolar lavage cells isolated from the lungs of severe COVID-19 patients [35,36], whilst other studies did not find an association between MIP-1β and severe disease [28,37].

The levels of IL-6, IL-8, and MIP-1β significantly decline at month 1 showing similar levels during the follow-up to those patients with mild/moderate disease. Overall, a similar dynamic was observed for IP-10, TNF-*α* and sCD25 during the study period. By contrast, the dynamic for MIP-1β, IL-1β, MIP-1*α* and IFN-*γ* showed a different evolution pattern with a significant increase at month 1 and/or 3 among patients with mild/moderate disease compared to those with severe COVID-19. No differences were observed in the levels of these cytokines 6 months after of the acute SARS-COV-2 infection between patients with mild/moderate vs. severe disease. Therefore, the unbalance in the cytokine levels between severe and mild/moderate COVID-19 patients seems to be restored after 6 months of SARS-CoV-2 infection. A previous study by Lucas et al., [38] also reported no differences at the IL-6, IL-8 and IP-10 levels between mild/moderate and severe outcomes after 20 days of follow-up.

This study presents some limitations. First, the relatively small size of the study population mainly represented by men as during the first wave of COVID-19 in our institution 86% of hospitalized patients were men. However, we were able to accomplish a very close follow-up of the clinical outcomes, viral and host immune factors in this population allowing interesting observations. Although we have identified IgG after 6 months of SARS-CoV-2 infection we did not assess their neutralization activity and therefore the risk upon new SARS-CoV-2 re-infections. External validation should be assessed to confirm the predictive value of our findings.

In conclusion, in a well-characterized cohort of hospitalized COVID-19 with mild/moderate and severe disease and close clinical follow-up during and after 6 months of SARS-CoV-2 infection we have recognized early host immune predictors of severity. Higher levels of IgA, IgM, and IgG and the specific cytokines IL-6, IL-8, and MIP-1β during acute infection were observed in those patients with a severe COVID-19 outcome. Conversely, higher levels of IL-1β and IFN-*γ* and at month 1 and MIP-1β, IL-1β, MIP-1*α* and IFN-*γ* month 3 were observed among patients with mild/moderate diseases compared to those with severe COVID-19. IgG against SARS-CoV-2 persisted after 6 months of the diagnosis.

## Acknowledgments

We would like to thank all the members of COHVID-GS and IISGS Biobank, patients, nursing staff and Celta Ingenieros (A Coruña, Spain) for their kind supply of ddPCR reagents.

## Members of COHVID-GS (Galicia Sur Health Research Institute)

Alejandro Araujo, Jorge Julio Cabrera, Víctor del Campo, Manuel Crespo, Alberto Fernández, Beatriz Gil de Araujo, Carlos Gómez, Virginia Leiro, María Rebeca Longueira, Ana López-Domínguez, José Ramón Lorenzo, María Marcos, Alexandre Pérez, María Teresa Pérez, Lucia Patiño, Sonia Pérez, Silvia Pérez-Fernández, Eva Poveda, Cristina Ramos, Benito Regueiro, Cristina Retresas, Tania Rivera, Olga Souto, Isabel Taboada, Susana Teijeira, María Torres, Vanesa Val, Irene Viéitez

