## Supplementary_Material for "Elevated anti-SARS-CoV-2 antibodies and IL-6, IL-8, MIP-1β, early predictors of severe COVID-19"

### Supplementary Methods

#### ddPCR quantification

The one-step RT-PCR reaction mixture was assembled from 1x supermix, 20U/µL reverse transcriptase, 15mM DTT, 1x 2019-nCoV CDC ddPCR Triplex Probe Assay, 5.5µL of RNA sample and RNase/DNase free water up to a final volume of 22µL. Then, the reaction mixture was converted to droplets with the QX200 droplet generator (Bio-Rad Laboratories). Droplet-partitioned samples were amplified in a C1000 Touch™ Thermal Cycler (Bio-Rad Laboratories) under these cycling conditions: 50°C for 60 min (reverse transcription), 95°C for 10 min (enzyme activation), followed by 40 cycles of 94°C for 30 s (denaturation) and 55°C for 1 min (annealing and extension), 98°C for 10 min (enzyme deactivation), and 4°C for 30 min (droplet stabilization). The cycled plate was read in the FAM and HEX channels using the QX200 Reader (Bio-Rad Laboratories). All experiments included a positive SARS-CoV-2 control (Exact Diagnostics SARS-CoV-2 Standard, Exact Diagnostic), a negative control (Exact Diagnostics SARS-CoV-2 Negative, Exact Diagnostic) and a no template control. All controls and samples are tested in duplicate. The data analysis was performed using the QuantaSoft Analysis Pro Software (Bio-Rad Laboratories). As quality control, a result was accepted if it had equal or more than 10.000 droplets. In addition, a sample was considered positive for SARS-CoV-2, and a quantitative data was reported, if it had both a minimal concentration of N (N1 and/or N2) of 0.1 copies/µL of 1X ddPCR reaction, and two or more positive droplets for N1 and/or N2 were present. For *RPP30*, the minimal concentration was up to 0.2 copies/µL of 1X ddPCR reaction and four or more positive droplets. Absolute quantification data was expressed in copies per milliliter of swab.

The serial dilutions of the RNA extracted from the commercial positive control for the experimental linearity study range from 200 copies/µL to 0.39 copies/µL for N1 and N2, and from 200 copies/µL to 0.78 copies/µL for *RPP30* (according to the minimal concentration established by the assay). The mean values obtained were compared with the expected values for each targets (N1, N2 and *RP30*), and the results were expressed in $\log_{10}$ (copies/reaction). Finally, we also normalized the viral load of each clinical sample referring the SARS-CoV-2 viral load quantification to the cellular quality of the swab (using the copy number of *RPP30* as reference for diploid cells), so the results were expressed in copies/$10^4$ cells for the viral load dynamic study. We also converted the SARS-CoV-2 viral load in $\log_{10}$ (copies/$10^4$ cells) in the graphics.

**Antibodies measurements**

The immunoassay plates (MaxiSorp, Nunc) were from Fisher Scientific and the SARS-CoV-2 nucleoprotein NP antigen from MyBiouSource (ref. MBS596190). The antibody standards IgA, IgM and IgG were from Invitrogen (human IgA ref. 31148; human IgM ref. 31146) and MP Biomedical (human IgG ref. 0855908). The enzyme conjugates were from Invitrogen (anti-human IgA-HRP ref. PA174395; anti-human IgM-HRP ref. 31415) and Sigma (anti-human IgG-HRP ref. A0170). Calibration curves were included in all plates to estimate the concentration of each antibody type in the samples. Serial dilutions of each antibody standard IgA, IgM or IgG were performed with 50 mM carbonate buffer pH 9.4 (dilution factor 1/4) in the range of 3.9 ng/mL – 16 µg/mL for IgA and IgG and 1.9 ng/mL – 8 µg/mL for IgM and were used to coat duplicate wells in parallel with the NP coating step for sample analysis. The serum samples were analyzed on NP-coated wells in triplicate, while duplicate control (non-coated) wells were also included to eliminate any signals resulting from non-specific binding of serum components to the wells. Calibration curves for each antibody type (IgA, IgM or IgG) were constructed by fitting the absorbance values (A450nm) to a sigmoidal 4-parameter logistical model using GraphPad Prism. The concentration of each antibody was then interpolated from its corresponding calibration curve using the corrected (A-A0) values for each sample, where A was the average absorbance of the NP-coated wells and A0 the average absorbance of the control (non-coated) wells. Pre-pandemic serum samples (n=30) were used as controls to set the background values of the in-house developed ELISA.

**Supplementary Figure 1.** Linear regression analysis of the RT-ddPCR SARS-CoV-2 assay.

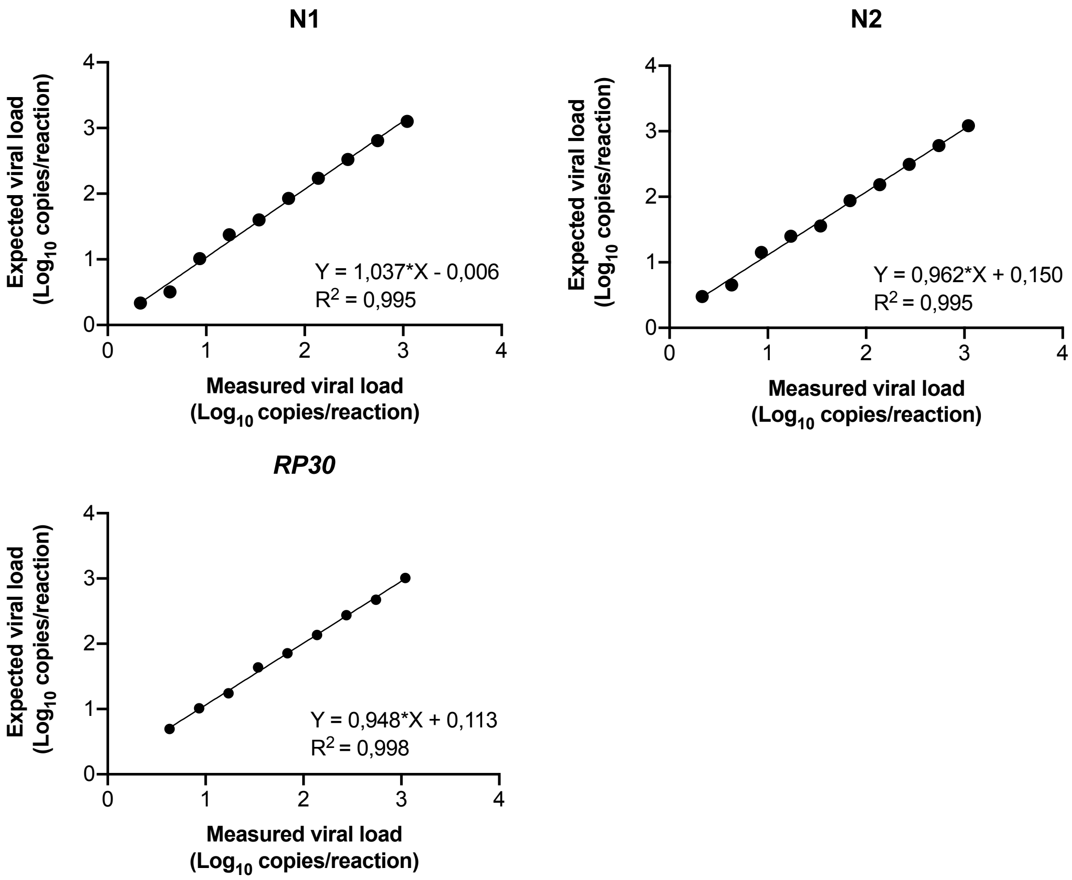


The graphics show the linearity between the expected and the observed values quantified from the serial dilutions of the SARS-CoV-2 commercial control (Exact Diagnostics SARS-CoV-2 Standard, Exact Diagnostic) and for the three targets included in the assay (N1, N2 and *RP30*). The quantification is represented as copies of target per reaction (in $Log_{10}$ format), and each black dot represents the mean of the replicates quantified for each dilution. Linear regression equation and correlation factor (r-square) are also included in the plots.

**Supplementary Table 1.** IgA, IgM, IgG levels at the different points of the study periods.

| | IgA (µg/mL) | IgM (µg/mL) | IgG (µg/mL) |
|---|---|---|---|
| **Baseline** | 6.72 (0.00 – 22.54) | 0.56 (0.12 – 1.30) | 26.67 (9.14 – 86.38) |
| **Day 3** | 18.00 (7.43 – 32.28) | 1.01 (0.54 – 2.43) | 70.81 (38.96 – 133.50) |
| **Day 6** | 26.25 (11.07 – 61.32) | 2.21 (0.61 – 4.52) | 109.30 (52.89 – 174.01) |
| **Month 1** | 2.45 (0.27 – 6.97) | 0.52 (0.08 – 0.88) | 89.12 (61.40 – 114.05) |
| **Month 3** | 0.00 (0.00 – 2.01) | 0.10 (0.00 – 0.53) | 63.70 (33.97 – 87.76) |
| **Month 6** | 0.00 (0.00 – 0.49) | 0.00 (0.00 – 0.15) | 15.93 (9.88 – 25.01) |

Data are represented as median (IQR)
n=18

**Supplementary Table 2.** Cytokine levels (severe vs. moderate)

| | Moderate patients | Severe patients | p-value |
|---|---|---|---|
| Baseline cytokines (pg/mL) | | | |
| IL-6 | 5.44 (2.93 – 11.63) | 32.72 (20.78 – 107.92) | **0.002** |
| IL-8 | 6.43 (4.65 – 9.51) | 19.99 (9.83 – 45.67) | **0.007** |
| IP-10 | 515.40 (141.30 – 633.13) | 1083.80 (439.15 – 4613.25) | 0.102 |
| TNF-$\alpha$ | 8.12 (7.31 – 11.56) | 11.55 (10.40 – 17.45) | 0.083 |
| sCD25 | 1221.50 (775.33 – 2806.00) | 2160.00 (1541.25 – 2877.00) | 0.250 |
| MIP-1β | 15.37 (8.99 – 17,51) | 25.29 (19.53 – 34.19) | **0.001** |
| IL-1β | 1.50 (1.31 – 2.02) | 1.65 (1.07 – 1.77) | 0.750 |
| MIP-1$\alpha$ | 12.74 (7.95 – 15.47) | 18.52 (5.02 – 24.21) | 0.682 |
| IFN-$\gamma$ | 35.91 (26.79 – 53.89) | 47.59 (36.60 – 101.63) | 0.213 |
| Month 1 cytokines (pg/mL) | | | |
| IL-6 | 2.40 (1.37 – 2.80) | 1.08 (0.88 – 8.94) | 0.635 |
| IL-8 | 4.73 (3.47 – 7.06) | 5.18 (3.04 – 6.13) | 0.945 |
| IP-10 | 67.57 (38.28 – 103.67) | 107.30 (97.54 – 114.33) | 0.106 |
| TNF-$\alpha$ | 7.75 (6.91 – 9.73) | 6.98 (5.02 – 10.13) | 0.635 |
| sCD25 | 824.75 (742.48 – 997.3) | 739.6 (201.06 – 1848.33) | 0.839 |
| MIP-1β | 12.60 (11.30 – 15.71) | 11.81 (7.08 – 25.45) | 0.839 |
| IL-1β | 1.83 (1.22 – 2.74) | 1.08 (0.90 – 1.44) | **0.024** |
| MIP-1$\alpha$ | 17.88 (13.28 – 24.16) | 14.48 (7.02 – 18.58) | 0.347 |
| IFN-$\gamma$ | 46.28 (41.57 – 57.21) | 25.10 (21.74 – 33.28) | **0.024** |
| Month 3 cytokines (pg/mL) | | | |
| IL-6 | 1.88 (0.84 – 2.81) | 4.94 (4.45 – 5.42) | 0.240 |
| IL-8 | 5.21 (2.97 – 8.05) | 2.03 (0.25 – 4.99) | 0.060 |
| IP-10 | 57.29 (24.32 – 131.25) | 71.34 (47.64 – 99.31) | 0.699 |
| TNF-$\alpha$ | 5.66 (4.72 – 6.70) | 4.94 (4.45 – 5.42) | 0.190 |
| sCD25 | 730.40 (549.55 – 944.25) | 782.5 (621.95 – 1092.50) | 0.797 |

| | | | |
|---|---|---|---|
| MIP-1β | 12.24 (7.35 – 18.75) | 5.27 (4.92 – 6.47) | **0.012** |
| IL-1β | 1.75 (1.14 – 1.99) | 0.26 (0.19 – 1.16) | **0.029** |
| MIP-1$\alpha$ | 15.46 (11.23 – 23.14) | 8.23 (0.15 – 10.70) | **0.007** |
| IFN-$\gamma$ | 50.80 (36.80 – 63.43) | 28.93 (22.74 – 33.62) | **0.029** |
| Month 6 cytokines (pg/mL) | | | |
| IL-6 | 2.44 (0.09 – 3.06) | 1.21 (0.09 – 2.69) | 0.660 |
| IL-8 | 2.53 (1.14 – 4.09) | 1.34 (0.69 – 7.74) | 0.859 |
| IP-10 | 42.97 (31.39 – 87.07) | 62.86 (52.44 – 84.92) | 0.256 |
| TNF-$\alpha$ | 6.15 (5.31 – 6.69) | 5.61 (5.24 – 6.92) | 0.733 |
| sCD25 | 533.00 (486.80 – 714.40) | 713.85 (535.00 – 989.75) | 0.216 |
| MIP-1β | 9.52 (6.39 – 16.89) | 8.54 (7.46 – 11.46) | 1.000 |
| IL-1β | 1.86 (0.63 – 2.43) | 0.78 (0.38 – 1.32) | 0.145 |
| MIP-1$\alpha$ | 32.38 (24.26 – 44.49) | 24.85 (8.46 – 25.39) | 0.069 |
| IFN-$\gamma$ | 60.12 (47.81 – 95.37) | 48.17 (38.02 – 55.31) | 0.098 |

Data are median (IQR)
Mann-Whitney U test was performed. P-values < 0,05 were considered significant and are highlighted in bold.
